# Validation of Serum Neurofilament Light Chain as a Biomarker of Parkinson’s Disease Progression

**DOI:** 10.1101/762237

**Authors:** Brit Mollenhauer, Mohammed Dakna, Tzu-Ying Liu, Douglas Galasko, Tatiana Foroud, Henrik Zetterberg, Sebastian Schade, Roland G. Gera, Wenting Wang, Feng Gao, Niels Kruse, Mark Frasier, Lana M. Chahine, Christopher S. Coffey, Andrew B. Singleton, Tanya Simuni, Daniel Weintraub, John Seibyl, Arthur W. Toga, Caroline M. Tanner, Karl Kieburtz, Kenneth Marek, Andrew Siderowf, Jesse M. Cedarbaum, Samantha J. Hutten, Claudia Trenkwalder, Danielle Graham

## Abstract

**Objective:** To assess neurofilament light chain (NfL), as a biomarker for Parkinson’s disease (PD).

**Methods:** We quantified NfL in (1) longitudinal CSF samples from PD, other cognate/neurodegenerative disorders (OND), and healthy controls (HC); (2) a cross-sectional cohort with paired CSF and serum samples from participants with PD, OND, and HC, and (3) a large longitudinal validation cohort with serum samples from PD, OND, HC, prodromal conditions, and mutation carriers.

**Results:** In the longitudinal discovery cohort (1) NfL in CSF was highest in OND and higher in PD vs. HC across all visits (p<0.05) but did not change longitudinally. In the cross-sectional cohort (2) paired CSF and serum NfL samples were highly correlated (Spearman’s rank 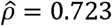; p<10^-6). In the large validation cohort (3) mean baseline serum NfL was higher in PD (13±7.2pg/ml) vs. HC (12±6.7pg/ml; p=0.0336) and was highest in OND (18±7pg/ml; p=0.0351). Serum NfL increased longitudinally in PD vs. HC (p<0.01). Longitudinal motor scores were positively longitudinally associated with NfL, whereas some cognitive scores showed a negative longitudinal association with NfL.

**Conclusions:** NfL levels in serum samples are increased in PD vs. HC, increase significantly over time, and correlate with clinical measures of PD severity. Although the specificity of NfL in PD is low and more specific biomarkers are needed, serum NfL is the first blood-based biomarker candidate that could support disease stratification (PD vs. OND), track clinical progression, and possibly assess responsiveness to neuroprotective treatments. NfL as a biomarker of response to neuroprotective interventions remains to be determined.

**Funding sources for study:** PPMI is sponsored by the Michael J. Fox Foundation for Parkinson’s Research (MJFF) and is co-funded by MJFF, Abbvie, Avid Radiopharmaceuticals, Biogen Idec, Bristol-Myers Squibb, Covance, Eli Lilly & Co., F. Hoffman-La Roche, Ltd., GE Healthcare, Genentech, GlaxoSmithKline, Lundbeck, Merck, MesoScale, Piramal, Pfizer and UCB. The funders had no role in the design and conduct of the study, in the collection, management, analysis, and interpretation of the data, in the preparation, review, or approval of the manuscript or in the decision to submit the manuscript for publication.

**Financial Disclosure/Conflict of Interest concerning the research related to the manuscript:** Brit Mollenhauer, Douglas Galasko, Tatiana Foroud, Lana M. Chahine, Christopher S. Coffey, Andrew B. Singleton, Tanya Simuni, Daniel Weintraub, John Seibyl, Arthur W. Toga, and Caroline M. Tanner received funding from The Michael J. Fox Foundation for Parkinson’s Research.

Mohammed Dakna, Tzu-Ying Liu, Henrik Zetterberg, Sebastian Schade, Roland G. Gera, Wenting Wang, Feng Gao, Niels Kruse, Mark Frasier, Jesse M. Cedarbaum, Samantha J. Hutten, Claudia Trenkwalder, and Danielle Graham report no disclosures.

## Introduction

Two major obstacles hamper the success of translational Parkinson’s disease (PD) research: 1) currently no longitudinal fluid biomarker for PD correlates with disease progression; 2) a definite diagnosis of PD can only be made by autopsy and the rate of clinical misdiagnoses, especially early in the disease, is reported to be high.^1, 2^ Therefore, biomarkers that could be used for diagnosis and as progression markers in PD are needed.

Neurofilaments are highly phosphorylated neuronal cytoskeleton components maintaining neuronal structure and determining axonal caliber. The 68 kDa neurofilament light chain (NfL) is released into extracellular fluids in response to axonal damage.^3, 4^ CSF NfL levels seem to reflect progression of various neurological conditions, including multiple sclerosis and neurodegenerative dementias.^5, 6^ NfL levels have been shown to differentiate sporadic PD from multiple system atrophy and progressive supranuclear palsy.^4, 7, 8^ While one small study did not show a longitudinal change in NfL levels during progression in PD,^9^ a recent single center study on plasma NfL showed a modest correlation of plasma NfL with the MDS-UPDRS III.^10^ Longitudinal analyses of NfL in larger multicenter cohorts, including prodromal and monogenetic participants, have not been carried out.

We hypothesized that NfL would: 1) be higher in PD than in healthy controls (HC), 2) be even greater in participants with other cognate or neurodegenerative disorders (OND), 3) increase over time with disease progression, 4) be higher in prodromal and asymptomatic mutation carriers than in controls, and 5) correlate with clinical measures and/or imaging indices of progression.

## Methods

### Study population

#### The DeNoPa- and Kassel-discovery cohorts

As previously described,^15,16^ DeNoPa (**de no**vo **Pa**rkinson’s disease) is an on-going prospective longitudinal, single center study. NfL measurements were obtained in CSF of 176 participants, including (1) newly diagnosed, drug-naïve PD patients, (2) age-, sex- and education-matched HC and (3) participants, who were initially enrolled as having PD but upon clinical follow-up had their diagnoses revised to cognate or neurodegenerative disorder (OND). In- and exclusion criteria for the longitudinal DeNoPa cohort including the identification of OND have been published previously.^11, 12^ The number of participants in each group is given in **table 1**. The clinical assessment battery of the core cohort was published previously.^13^ In brief, motor assessment used the revised Unified Parkinson’s Disease Rating Scale published by the Movement Disorder Society (MDS-UPDRS III and total score).^14^ Cognition was assessed by Mini Mental Status Examination (MMSE). The pre-specified time points for CSF collection were baseline, 6-, 24-, 48-, and 72-month visits (**table 1**).

**Table 1a:** Demographics and NfL measures in CSF of the longitudinal discovery cohort (DeNoPa-cohort)

| Parameter | Level | Healthy Control (HC) | Parkinson's Disease (PD) | Other Neurodegenerative Diseases (OND) | FDR adjusted p-values |
| --- | --- | --- | --- | --- | --- |
| **N** |  | 61 | 98 | 17 |  |
| **Sex** |  |  |  |  |  |
|  | Female | 20 (32.8%) | 33 (33.7%) | 4 (23.5%) | 0.81<sup>1</sup> |
|  | Male | 41 (67.2%) | 65 (66.3%) | 13 (76.5%) |  |
| **Age at baseline** |  |  |  |  |  |
|  | mean±SD | 65 ± 6.8 | 65 ± 9.9 | 67 ± 7.6 | 0.81<sup>2</sup> |
|  | median (min; max) | 66 (44; 84) | 66 (40; 84) | 66 (53; 78) |  |
| **MDS-UPDRS part III** |  |  |  |  |  |
|  | mean±SD | 0.62 ± 1.4 | 23 ± 11 | 26 ± 13 | < 0.01<sup>2</sup> |
|  | median (min; max) | 0 (0; 6) | 22 (3; 54) | 24 (6; 48) |  |
| **MDS-UPDRS total score** |  |  |  |  |  |
|  | mean±SD | 3.3 ± 3.3 | 36 ± 17 | 43 ± 19 | < 0.01<sup>2</sup> |
|  | median (min; max) | 2 (0; 15) | 36 (7; 84) | 43 (14; 74) |  |
| **MMSE total score** |  |  |  |  |  |
|  | mean±SD | 29 ± 1.2 | 28 ± 1.4 | 27 ± 2.4 | 0.05<sup>1</sup> |
|  | median (min; max) | 29 (26; 30) | 29 (22; 30) | 28 (21; 30) |  |
| **Baseline** |  |  |  |  |  |
| N |  | 61 | 98 | 17 |  |
| **CSF NfL [pg/ml]** |  |  |  |  |  |
|  | mean±SD | 543 ± 250 | 675 ± 384 | 1063 ± 670 | < 0.01<sup>2</sup> |
|  | median (min; max) | 494<br>(249; 1626) | 562<br>(170; 2477) | 830<br>(416; 2876) |  |
|  | Missing | 9 | 19 | 0 |  |
| **24 months Follow-up** |  |  |  |  |  |
| N |  | 60 | 88 | 18 |  |
| **CSF NfL [pg/ml]** |  |  |  |  |  |
|  | mean±SD | 566 ± 261 | 1170 ± 2486 | 1240 ± 987 | < 0.01<sup>2</sup> |
|  | median (min; max) | 515<br>(259; 1618) | 669<br>(169; 16449) | 887<br>(519; 3878) |  |
|  | Missing | 28 | 45 | 6 |  |
| **48 months Follow-up** |  |  |  |  |  |
| N |  | 56 | 85 | 17 |  |
| **CSF NfL [pg/ml]** |  |  |  |  |  |
|  | mean±SD | 611 ± 290 | 751 ± 363 | 1025 ± 476 | 0.02<sup>2</sup> |
|  | median (min; max) | 531<br>(271; 1630) | 688<br>(201; 1869) | 984<br>(426; 1677) |  |
|  | Missing | 29 | 42 | 10 |  |
| **72 Months Follow-up** |  |  |  |  |  |
| N |  | 53 | 76 | 17 |  |
| **CSF NfL [pg/ml]** |  |  |  |  |  |
|  | mean±SD | 643 ± 228 | 977 ± 903 | 1016 ± 532 | 0.03<sup>2</sup> |
|  | median (min; max) | 605<br>(395; 1384) | 762<br>(295; 5298) | 888<br>(582; 1938) |  |
|  | Missing | 32 | 48 | 12 |  |

<sup>1</sup> Pearson's Chi-squared test
<sup>2</sup> Kruskal-Wallis rank sum test
**Abbreviations:** Movement Disorders Society Unified Parkinson's Disease rating Scale (MDS-UPDRS); Neurofilament light chain (NfL); Standard deviation (SD)
\* At 24 months follow-up all participants were reassessed as to the differential diagnosis (by clinical history, response to levodopa, and neurological examination) 17 returning disease participants were diagnosed with a different disease including atypical PD (as published<sup>12</sup>): PSP (n=3); MSA (n=1), Dementia with Lewy Bodies (DLB; n=2), one each with dystonic and cerebellar tremor and in nine cases the diagnosis remained unclear but suggested a progressive neurodegenerative disorder. This group was separated from the PD group and classified as other neurodegenerative disorders (OND).

The second discovery cohort included paired CSF and serum samples from the cross-sectional Kassel-cohort and comprised 150 PD and 344 OND (**table 3**). We also added 20 randomly selected HC from the DeNoPa cohort who were already included in the first discovery cohort. CSF and serum samples from the cross-sectional trainings cohort were collected from in-patients with careful clinical phenotyping that included magnetic resonance imaging (MRI) to determine structural abnormalities, quantitative levodopa testing as published^15^, smell identification test, MMSE followed by further cognitive testing and video-supported polysomnography (vPSG) to determine REM sleep behavior disorder (RBD) in a subset of patients. The phenotyping was undertaken in accordance with established criteria for PD, multiple system atrophy (MSA), dementia with Lewy bodies (DLB), progressive supranuclear palsy (PSP), corticobasal degeneration (CBD) and frontotemporal dementia (FTD). Participants with marked vascular lesions in 1.5 tesla magnetic resonance imaging (MRI) indicative for a vascular comorbidity and participants with normal pressure hydrocephalus by MRI were excluded.

#### The PPMI validation cohort

As previously described,^5,9^ PPMI (Parkinson’s Progression Marker Initiative) is an on-going prospective longitudinal, observational, international multicenter study that aims to identify PD biomarkers. We retrieved the clinical data from the PPMI data portal at the Laboratory of Neuroimaging (LONI) at the University of Southern California on June 6th, 2019 (www.ppmi-info.org) and obtained all the available aliquots at prespecified time points from the PPMI population. This generated serum NfL measurements of 1190 participants, including (1) newly diagnosed, drug-naïve PD patients, (2) age- and sex-matched HC, (3) prodromal participants with RBD or hyposmia, (4) the PPMI genetic cohort comprised individuals with pathogenic gene mutations in LRRK2, GBA, or SNCA, and (5) participants initially enrolled as PD but who had their diagnoses revised to OND upon clinical follow-up. The number of participants in each group and the demographics and clinical data are in **table 4**.

In PPMI the inclusion criteria for PD participants were the following: (1) aged over 30 years; (2) presence of two of the following: bradykinesia, rigidity and resting tremor, or an asymmetric resting tremor, or asymmetric bradykinesia; (3) diagnosis made within the last 24 months; (4) PD drug-naivety, and (5) dopamine transporter deficit in the putamen on the 123-I Ioflupane dopamine transporter imaging (DaT) by central reading. Isolated RBD participants met the following criteria: (1) men or women aged over 60 years, and (2) confirmation of RBD by video supported polysomnography (vPSG) with central reading and/or clinical diagnosis of RBD by site investigator including existing PSG as described. The central vPSG interpretation was based on the following criteria: (i) 18% of any EMG activity in m. mentalis, 32% of any EMG activity in mentalis and flexor digitorum superficialis (FDS) (in 3 s bins), (ii) 27% of any EMG activity in m. mentalis, 32% of any EMG activity in m. mentalis and FDS (in 30 s bins). In two cases a central PSG reading was not available due to technical difficulties with electronic PSG transfer, but these participants had a clinical diagnosis of isolated RBD (iRBD) by site investigator including prior PSG and also had to show decreased dopamine transporter imaging. Hyposmic participants were 60 years or older with olfaction at or below the 10th percentile by age and sex as determined by the University of Pennsylvania Smell Identification Test (UPSIT). All participants with iRBD and hyposmic participants also required confirmation from the imaging core at the Institute for Neurodegenerative Disorders (IND) that screening by DaT (or V-MAT-2-PET scan for sites where DaT is not available) was read as eligible. About 80% of the prodromal participants were selected with DaT deficit similar to participants with early PD, and 20% were selected with no DaT deficit. Prodromal participants without DaT deficit were similar in age, sex, and risk profile to those with mild to moderate dopamine transporter deficit. Exclusion criteria can be found in the study protocol at http://www.ppmi-info.org/study-design/research-documents-and-sops/.

Participants with the following mutations were included in the analysis of the genetic cohort: SNCA rs356181, SNCA rs3910105, LRRK2 rs34778348, GBA N370S rs76763715, SREBF1 rs11868035, LRRK2 rs34637584, MAPT rs17649553, SNCA rs3910105, SNCA rs356181, LRRK2 rs34637584, LRRK2 rs35801418, LRRK2 rs35870237, LRRK2 rs34995376. The prespecified time points for aliquot collection were baseline, 6-, 12-, 24-, 36-, 48- and 60-month visits for PD and HC, and baseline, 6-,12-, 24- and 36-month visits for prodromal and monogenetic participants. For the genetic cohort unaffected GBA mutation carriers the available follow-up was until the 24-month visit.

The clinical assessment battery, mainly of the core cohort, is described on the PPMI website and has been published previously.^13^ In brief, motor assessment used the MDS-UPDRS III and total score. Use of medications for PD was recorded at each visit after baseline assessment and is expressed as levodopa equivalent daily doses (LEDD).^16^

Cognitive testing included the Montreal Cognitive Assessment (MoCA) and psychometric tests of memory [Hopkins Verbal Learning Tests (HVLT) with immediate/total recall (HVLT-IR), discrimination recognition (HVLT-DG) and retention (HVLT-RT)], processing speed/attention [Symbol Digit Modality Test (SDMT)], executive function/working memory [WMS-III Letter-Number Sequencing Test (LNS)] and visuospatial abilities [Benton Judgment of Line Orientation test (BJLO)].^17^

Dopamine SPECT imaging was performed by DaT using standardized methods.^9^ Quantitative DaT measures of Striatal Binding Ratio (SBR) of caudate, putamen, or striatal uptake were used in our analyses.

#### Sample processing and immunoassays for NfL quantification

CSF samples of the DeNoPa-cohort were quantified by the commercial enzyme linked immunosorbent assay (ELISA from Uman-Diagnostics NfL ELISA kit, Umeå, Sweden).^18^ The assay was validated by 20 clinical CSF samples for inter-laboratory accuracy analysis in six different European laboratories.^19^ All other samples were analyzed by the newly developed Simoa NF-L Kit for the serum- and Simoa Neurology 4-plex A kit for the CSF-samples (in the cross-sectional discovery cohort) with the fully automated SIMOA® HD-1 analyzer (Quanterix, Lexington, MA, USA). NfL in the cross-sectional training cohort was quantified by the NF-light® assays from Quanterix (Lexington, USA): Simoa NF-L Beta Kit for serum and Simoa Neuro 4-plex A kit (multiplexing NfL, with the same antibodies as the single plex, GFAP, tau-protein and UCHL-1) for CSF with the fully automat SIMOA® HD-1 analyzer in duplicate with the investigator being blinded to the diagnosis. These assays have been shown to be up to 1,000-fold more sensitive than conventional immunoassays with limits of quantification in lower fg/ml range (important also for serum NfL measurements).^20^ For validation purposes we analyzed four CSF samples and three serum samples, diluted 1 in 100 to 1 in 800 and 1 in 4 to 1 in 32, respectively. In addition, recovery rates were determined in two spiked and non-spiked CSF and serum samples. NfL was detectable in all CSF and serum samples. Parallelism was observed for all CSF dilutions except for one sample diluted 1 in 800. For serum samples we observed parallelism for all samples except for one sample diluted 1 in 16 and one sample diluted 1 in 32. Recovery rates were more than 90% for CSF samples. Probably due to matrix effects, recovery rates were below 50% for serum samples. Here small amounts of the interphase generated during sample centrifugation were probably transferred to the assay plate. A direct comparison of CSF and serum samples on the single- and the multiplex assays showed that the measurements in single-versus multiplex assay are very comparable in levels. The NF-light® assay (Simoa NF-Light Advantage Kit; Quanterix, Lexington, USA) was used for the serum samples of the PPMI cohort diluted 1:4 and analyzed in duplicates at Quanterix (Lexington, MA, USA) with the investigators and analysts being blinded to the diagnosis. Three sets of endogenous quality controls were run on all plates. Three sets of spiked quality controls (QCs; recombinant neurofilament spiked into sample matrix) were prepared in bulk and run on all plates. Analytical assay parameters (i.e. quality control sample acceptance ranges) were determined prior to sample testing.

For the training cohorts all samples were collected in the morning under fasting conditions using Monovette® tubes (Sarstedt, Nümbrecht, Germany) for serum collection by venous puncture. CSF was collected in polypropylene tubes (Sarstedt, Nümbrecht, Germany) directly after the serum collection by lumbar puncture in the sitting position, tubes were centrifuged at 2500 g at room temperature (20 °C) for 10 minutes and aliquoted and frozen within 30 minutes after collection at −80°C until analysis. Before centrifugation white and red blood cell count in CSF was determined manually.^11, 21^

In the validation cohort (PPMI) serum was collected using standardized venous puncture procedures. Sample handling, shipment and storage were carried out according to the PPMI biologics manual (http://ppmi-info.org). Aliquots of 0.5 ml frozen serum were used for analysis.

#### Standard protocol approvals, registrations, and patient consent

Approval was received from the local ethical standards committee on human experimentation for all human participants. Written informed consent for research was obtained from all study participants. DeNoPa is registered in the German Register for Clinical trials (DRKS00000540), PPMI in clinicaltrials.gov as NCT01141023.

### Statistical analysis

#### Analysis of the discovery cohorts

The analysis of the longitudinal discovery cohort was carried out in congruency to PPMI described in more detail below with linear mixed model with log2NfL as response to longitudinal time and diagnostic groups. Since the longitudinal discovery cohort (DeNoPa) was age- and sex-matched no adjustment was carried out. The Kruskal-Wallis test was performed to find differences between more than three groups followed by pairwise comparisons using Mann-Whitney U tests adjusted using Tukey’s Honestly Significant Difference (HSD). The correlation of NfL in paired CSF and serum samples and the clinical measures in the discovery and validation cohort was analyzed by Spearman’s rank test.

#### Analysis of the validation cohort

Unlike the DeNoPa cohort, the PPMI cohort was not age- and sex-matched. We explored the association between log2NfL, age and sex among the HC by locally estimated scatterplot smoothing (LOESS) and a linear regression model. We used log transformation as the distribution of NfL was right skewed. To visualize the changes of NfL over time among different diagnostic groups, age- and sex-adjusted log2NfL was computed and plotted against the disease duration for patients with PD or OND and the time in the study for HC, unaffected mutation carriers, and prodromal participants. We compared the median baseline NfL using a linear regression model. To compare changes over time, we fitted a linear mixed model for log2NfL with disease duration, diagnostic groups and their interactions as covariates, adjusted by age and sex. For HC, unaffected mutation carriers, and prodromal participants, their disease durations were replaced by the time in the study.

Finally, we examined the association between the clinical measurements and NfL among patients with PD by fitting a linear median mixed model for each clinical measurement, with adjusted log2NfL, age, sex, disease duration, and whether the patient was on- or off-medication at the visit as the covariates.

The linear mixed models were fitted using the R package “lme4” and the linear median mixed model was fitted using the R package “lqmm”. For all cohorts the p-values were adjusted via controlling of the false discovery rate (FDR) using the Benjamini & Hochberg method. ^22^

## RESULTS

### NfL measures in the discovery cohorts (DeNoPa- and Kassel-cohorts)

Quantification of NfL was first carried out in CSF samples from the longitudinal DeNoPa-cohort. **Table 1** shows the baseline values of CSF NfL and clinical scores. Among the 176 participants, patients with OND had the highest baseline median CSF NfL as 839 pg/ml, followed by PD patients (562 pg/ml) and HC (494 pg/ml, p=0.01; **figure 1a**). Linear mixed effect model of log2NfL on CSF NfL diagnoses, time and their interactions did not show significant differences across the groups (**table 2**).

**Figure 1a:**
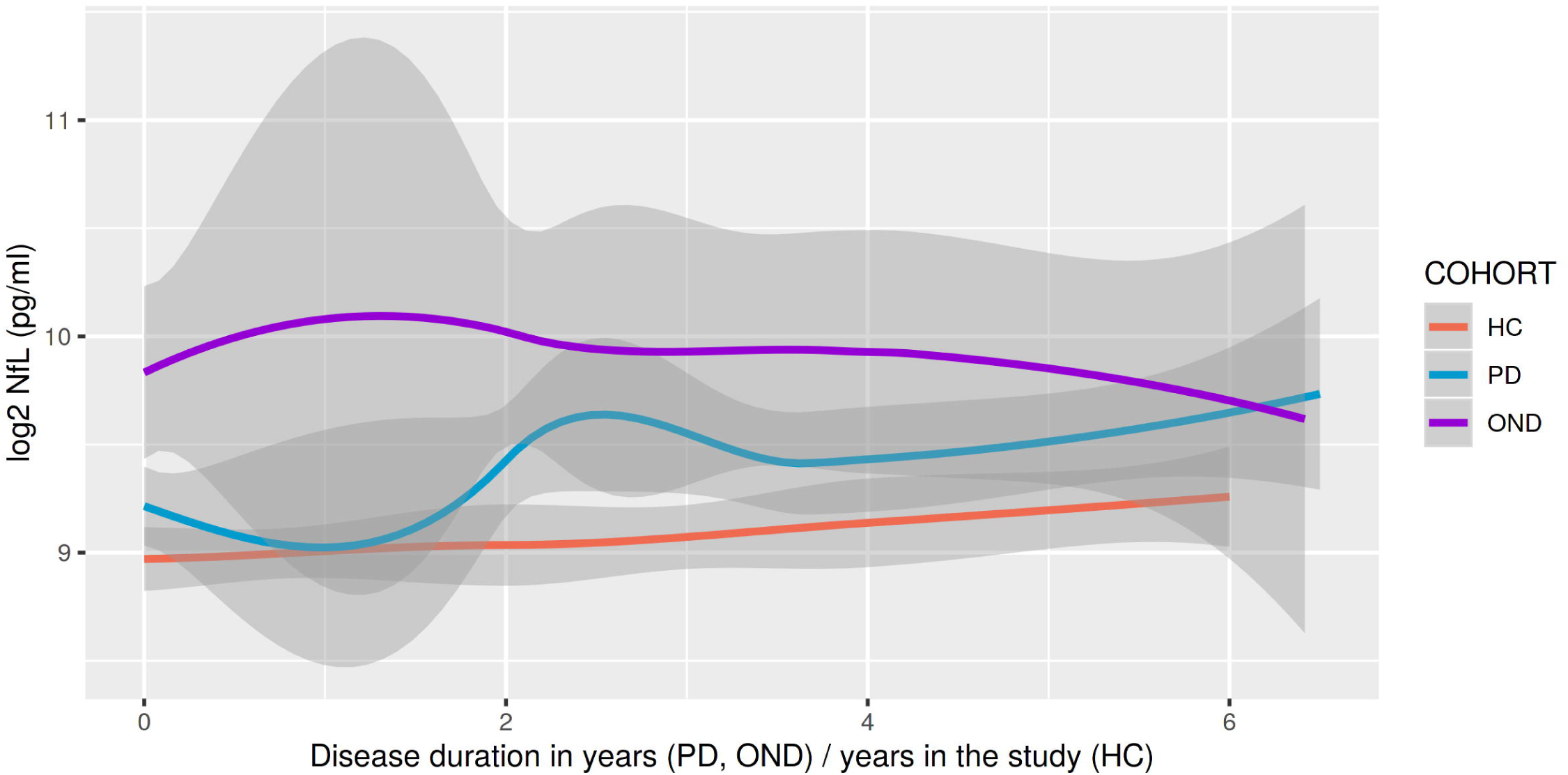
Log2-transformed CSF NfL levels at each visit in healthy controls (HC), Parkinson’s disease (PD), other neurodegenerative disorders (OND), in the longitudinal discovery cohort DeNoPa (demographics in table 1). The gray ribbon provides estimates of the standard error.

**Table 2:** Linear mixed effect model of log2NfL in CSF on diagnoses, time and their interactions, in the longitudinal discovery cohort (DeNoPa)

| Variable | Estimate | 2^Estimate | Std. Error | p-value |
| --- | --- | --- | --- | --- |
| (Intercept) | 6.3500 | 81.5713 | 0.3669 | <0.0001 |
| Age | 0.0366 | 1.0257 | 0.0053 | <0.0001 |
| Sex: male | 0.3058 | 1.2361 | 0.0993 | 0.0024 |
| Diagnostic group: Parkinson's Disease (PD) | 0.3292 | 1.2563 | 0.1036 | 0.0018 |
| Diagnostic group: Other Neurodegenerative Diseases (OND) | 0.8064 | 1.7489 | 0.1700 | <0.0001 |
| Time | 0.0189 | 1.0132 | 0.0168 | 0.2615 |
| Diagnostic group: Parkinson's Disease*TIME | 0.0261 | 1.0182 | 0.0209 | 0.2142 |
| Diagnostic group: Other Neurodegenerative Diseases*TIME | -0.0350 | 0.9760 | 0.0336 | 0.2986 |

In a second exploratory step, we analyzed a cross-sectional cohort where CSF and serum NfL levels were measured; the cohort demographics are reported in **table 3.** Among the 514 participants, patients with OND had the highest median CSF NfL (1895 pg/ml in CSF and 33pg/ml in serum, followed by PD patients (1438 pg/ml in CSF and 29 in serum pg/ml) and healthy participants (752 pg/ml in CSF and 15 in serum pg/ml, p<0.01). The strong correlation between CSF and serum NfL (by Spearman’s rank 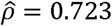; p < 10^-06; **figure 1b**) encouraged us to move to the validation step using serum samples from the longitudinal PPMI-cohort.

**Table 3:** Demographics and NfL measures in CSF and serum in the cross-sectional training cohort (Kassel-cohort)

| Parameter | Level | Healthy Control (HC) | Parkinson's Disease (PD) | Other Neurodegenerative Diseases (OND) | FDR adjusted p-value |
| --- | --- | --- | --- | --- | --- |
| **N** |  | 20 | 150 | 344 |  |
| **Sex** | Male | 14 (70.0%) | 100 (66.7%) | 224 (65.1%) | 0.91<sup>1</sup> |
|  | Female | 6 (30.0%) | 50 (33.3%) | 120 (34.9%) |  |
|  | Missing | 0 | 0 | 0 |  |
| **Age** | mean±SD | 69 ± 6.4 | 69 ± 9.6 | 71 ± 8.7 | 0.09<sup>2</sup> |
|  | median (min; max) | 70 (57; 86) | 71 (40; 86) | 72 (34; 88) |  |
|  | missing | 0 | 0 | 1 |  |
| **UPDRS part III** | mean±SD | 1.7 ± 2.2 | 31 ± 15 | 30 ± 14 | < 0.01<sup>2</sup> |
|  | median (min; max) | 1 (0; 7) | 30 (0; 75) | 28 (1; 83) |  |
|  | missing | 0 | 13 | 84 |  |
| **UPDRS total score** | mean±SD | 4.2 ± 5.9 | 55 ± 24 | 53 ± 23 | < 0.01<sup>2</sup> |
|  | median (min; max) | 3 (0; 25) | 54 (7; 135) | 50 (5; 135) |  |
|  | missing | 0 | 35 | 175 |  |
| **CSF NfL [pg/ml]** | mean±SD | 922 ± 446 | 2124 ± 2180 | 2779 ± 3544 | < 0.01<sup>2</sup> |
|  | median (min; max) | 752 (374; 2267) | 1438 (342; 15304) | 1895 (346; 35742) |  |
|  | missing | 0 | 1 | 3 |  |
| **Serum NfL [pg/ml]** | mean±SD | 18 ± 8.4 | 55 ± 105 | 44 ± 38 | < 0.01<sup>2</sup> |
|  | median (min; max) | 15 (5.8; 43) | 29 (1.5; 815) | 33 (5.7; 304) |  |
|  | missing | 0 | 1 | 3 |  |

<sup>1</sup> Fisher's Exact Test for count data
<sup>2</sup> Kruskal-Wallis rank sum test
**Abbreviations:** Unified Parkinson's Disease rating Scale (UPDRS); Neurofilament light chain (NfL); Cerebrospinal fluid (CSF)

**Figure 1b:**
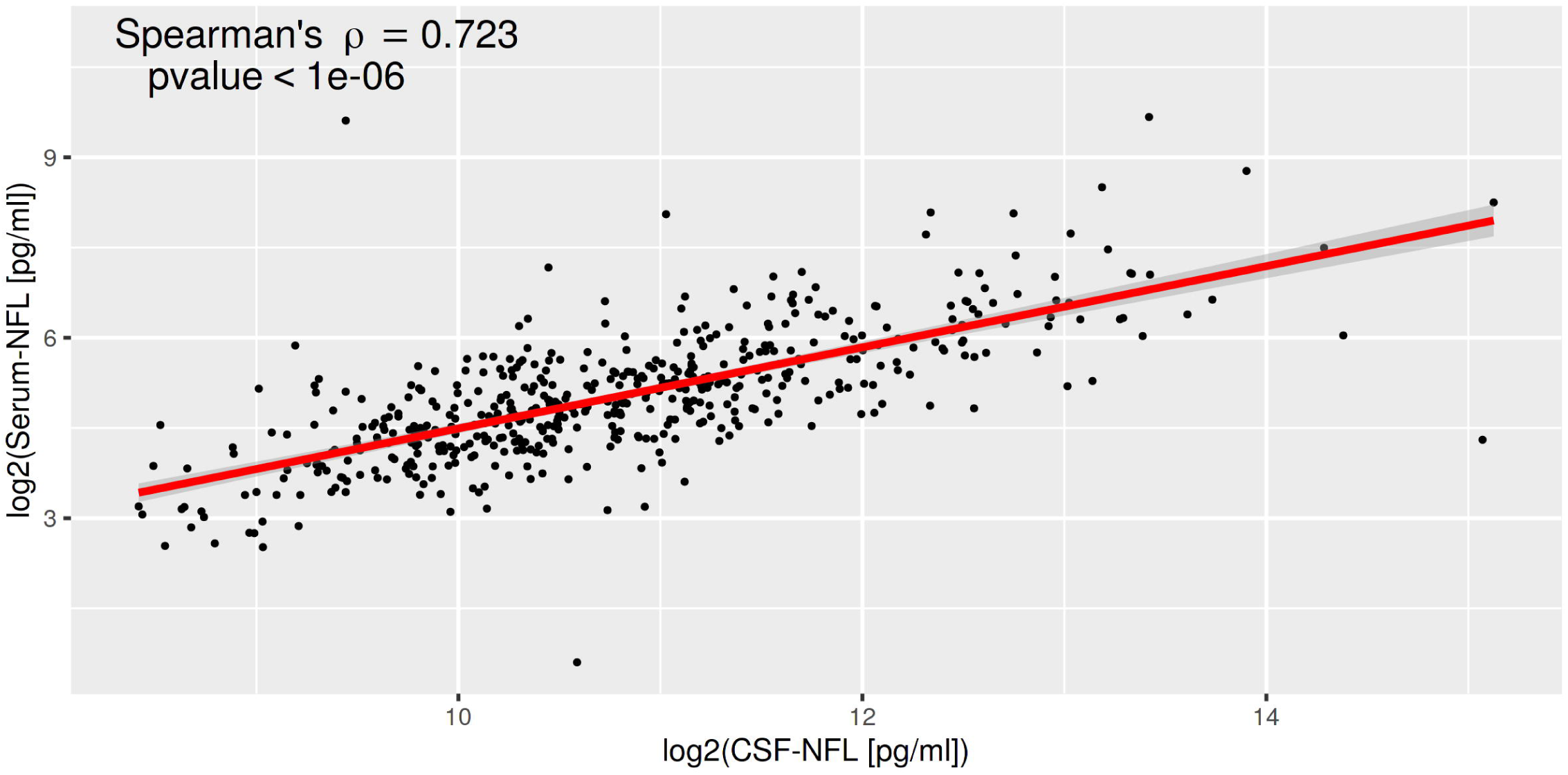
Spearman’s correlation of CSF and serum NfL in the cross-sectional discovery cohort (Kassel-cohort; demographics are in table 3)

**Figure 1c:**
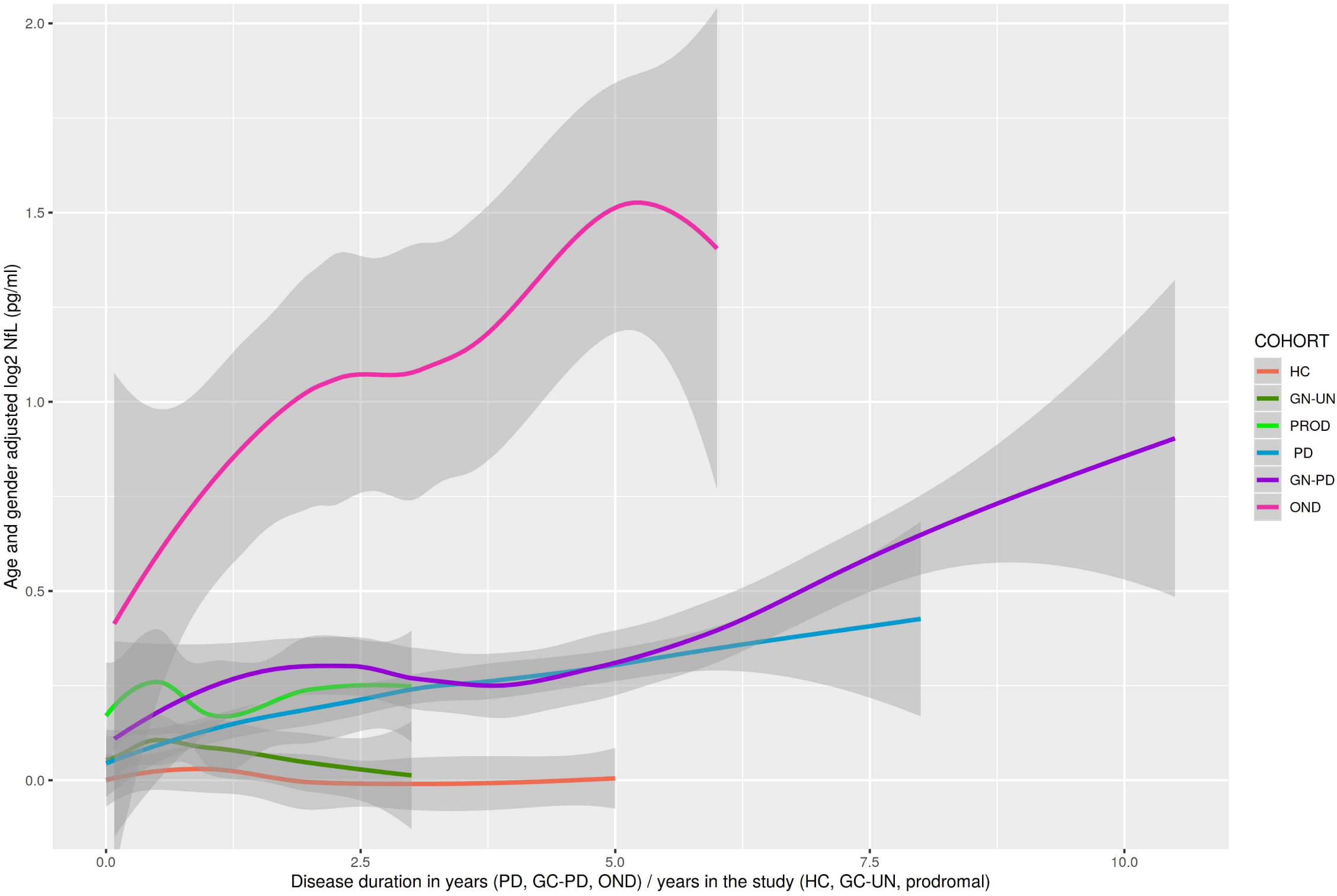
Age- and sex-adjusted log2 transformed serum NfL levels at each visit in healthy controls (HC), Parkinson’s disease (PD), other neurodegenerative disorders (OND), prodromal with hyposmia or isolated REM sleep behavior disorder (PROD) and the genetic cohort with affected- (GC-PD) and unaffected mutation carriers (GC-UN) in the PPMI-validation cohort (demographics in table 4). The gray ribbon provides estimates of the standard error.

### Serum NfL measures in the PPMI-validation cohort

The baseline characteristics of the PPMI-cohort (**table 4**) show significant differences in age and sex across the cohorts. Within the HC group there is an age- and sex-dependent change in serum NfL with a linear regression of log2NfL with age and sex as covariates explained 51% of the NfL variability (R^2^=0.51), which has also been shown in other cohorts.^23^ The downstream analyses of the PPMI data were all age- and sex-adjusted. Conditional on the median age of 63 years old and male sex, patients with OND had the highest baseline age- and sex-adjusted median serum NfL (16.23 pg/ml), followed by patients with genetic PD (13.36 pg/ml), prodromal participants (12.20 pg/ml), PD patients (11.73 pg/ml), unaffected mutation carriers (11.63 pg/ml) and HC (11.05 pg/ml, the F test of any differences among the medians had p-value<0.0001). Using a linear mixed model (**table 5**), median NfL increased by 3.35% per year of age (p<0.0001) and women had a median serum NfL level 6.79% higher than men (p=0.0002). Compared with the baseline median serum NfL of HC, the baseline median serum NfL of prodromal participants was 11.83% higher (p=0.039); the median serum NfL level at disease diagnosis for patients with genetic PD was 14.60% higher (p=0.0025); and the median serum NfL level at disease diagnosis for patients with OND was 42.15% higher (p=0.0086). The estimated yearly increase of serum NfL after disease diagnosis was 3.8% higher in PD patients compared with HC (p<0.0001), 2.5% higher in genetic PD patients (p=0.0175), and 16.19 % higher in patients with OND (p<0.0001).

**Table 4:** Demographics, clinical, dopamine transporter imaging and baseline serum NfL data in the PPMI validation cohort

| Parameter | Level | Healthy Control (HC) | Parkinson's Disease (PD) | Other Neurodegenerative Diseases (OND) | Prodromal (PROD) | Genetic Cohort – PD (GN-PD) | Genetic Cohort – Unaffected (GN-UN) | FDR-adjusted p-value |
| --- | --- | --- | --- | --- | --- | --- | --- | --- |
| **N** |  | 187 | 397 | 8 | 63 | 226 | 309 |  |
| **Sex** | female | 67 (35.8%) | 140 (35.3%) | 2 (25.0%) | 14 (22.2%) | 117 (51.8%) | 183 (59.2%) | < 0.01<sup>1</sup> |
|  | male | 120 (64.2%) | 257 (64.7%) | 6 (75.0%) | 49 (77.8%) | 109 (48.2%) | 126 (40.8%) |  |
| **Age** | mean±SD | 61 ± 11 | 62 ± 9.9 | 64 ± 6.8 | 69 ± 5.8 | 62 ± 10 | 62 ± 7.4 | < 0.01<sup>2</sup> |
|  | median (min; max) | 62 (31; 84) | 62 (34; 85) | 66 (53; 73) | 67 (59; 83) | 65 (32; 85) | 62 (34; 84) |  |
| **MDS-UPDRS part III** | mean±SD | 1.2 ± 2.2 | 21 ± 8.9 | 27 ± 5.4 | 3.9 ± 3.8 | 23 ± 11 | 2.7 ± 3.9 | < 0.01<sup>2</sup> |
|  | median (min; max) | 0 (0; 13) | 20 (4; 51) | 28 (18; 36) | 3 (0; 15) | 21 (4; 56) | 1 (0; 27) |  |
| **MDS-UPDRS Total Score** | mean±SD | 4.6 ± 4.4 | 32 ± 13 | 47 ± 10 | 12 ± 7.8 | 38 ± 18 | 8.6 ± 7.7 | < 0.01<sup>2</sup> |
|  | median (min; max) | 3 (0; 20) | 31 (7; 70) | 44 (37; 68) | 11 (0; 31) | 36 (7; 113) | 7 (0; 45) |  |
| **MoCA** | mean±SD | 28 ± 1.1 | 27 ± 2.3 | 27 ± 2.3 | 26 ± 3.5 | 26 ± 3.5 | 27 ± 2.3 | < 0.01<sup>2</sup> |
|  | median (min; max) | 28 (27; 30) | 28 (17; 30) | 28 (23; 30) | 27 (11; 30) | 27 (11; 30) | 27 (16; 30) |  |
| **HVLT-IR** | mean±SD | 26 ± 4.5 | 24 ± 5 | 26 ± 4.6 | 22 ± 5.3 | 24 ± 5.6 | 27 ± 4.9 | < 0.01<sup>2</sup> |
|  | median (min; max) | 26 (15; 35) | 25 (9; 36) | 25 (18; 32) | 21 (9; 33) | 24 (7; 34) | 27 (9; 36) |  |
| **HVLT-DG** | mean±SD | 10 ± 2.8 | 9.7 ± 2.6 | 9.9 ± 1.7 | 9.4 ± 2.2 | 9.9 ± 2.3 | 11 ± 1.8 | < 0.01<sup>2</sup> |
|  | median (min; max) | 11 (-4; 12) | 10 (-4; 12) | 10 (7; 12) | 10 (2; 12) | 10 (-1; 12) | 11 (0; 12) |  |
| **HVLT-RT** | mean±SD | 0.9 ± 0.18 | 0.85 ± 0.2 | 0.86 ± 0.17 | 0.77 ± 0.29 | 0.8 ± 0.25 | 0.88 ± 0.17 | < 0.01<sup>2</sup> |
|  | median (min; max) | 0.92 (0.27; 1.5) | 0.9 (0; 1.3) | 0.8 (0.71; 1.2) | 0.83 (0; 1.2) | 0.89 (0; 1.4) | 0.92 (0.12; 1.4) |  |
| **SDMT** | mean±SD | 47 ± 11 | 41 ± 9.8 | 38 ± 7.6 | 36 ± 11 | 38 ± 12 | 46 ± 9.3 | < 0.01<sup>2</sup> |
|  | median (min; max) | 46 (20; 83) | 42 (7; 82) | 40 (23; 46) | 35 (15; 56) | 39 (6; 75) | 47 (14; 74) |  |
| **LNS** | mean±SD | 11 ± 2.6 | 11 ± 2.7 | 10 ± 1.4 | 9.4 ± 2.9 | 9.6 ± 3.1 | 11 ± 2.8 | < 0.01<sup>2</sup> |
|  | median (min; max) | 11 (2; 20) | 11 (2; 20) | 10 (9; 13) | 9 (3; 17) | 10 (2; 18) | 11 (0; 20) |  |
| BJLOS | mean±SD | 13 ± 2 | 13 ± 2.1 | 13 ± 2.1 | 12 ± 2.3 | 12 ± 3 | 13 ± 2.1 | < 0.01<sup>2</sup> |
|  | median (min; max) | 14 (4; 15) | 13 (5; 15) | 13 (8; 15) | 12 (3; 15) | 12 (0; 15) | 13 (1; 15) |  |
| DaT-Scan PUTAMEN (SBR) | mean±SD | 2.1 ± 0.54 | 0.82 ± 0.28 | 0.63 ± 0.32 | NA | 0.76 ± 0.35 | NA | < 0.01<sup>2</sup> |
|  | median (min; max) | 2.1 (0.64; 3.9) | 0.79 (0.24; 2.2) | 0.55 (0.31; 1.3) | NA | 0.7 (0.21; 2.4) | NA |  |
| DaT-Scan CAUDATE (SBR) | mean±SD | 3 ± 0.6 | 2 ± 0.54 | 1.4 ± 0.64 | NA | 1.8 ± 0.6 | NA | < 0.01<sup>2</sup> |
|  | median (min; max) | 2.9 (1.3; 4.8) | 2 (0.39; 3.7) | 1.2 (0.74; 2.6) | NA | 1.8 (0.43; 3.9) | NA |  |
| DaT-Scan STRIATUM (SBR) | mean±SD | 2.6 ± 0.55 | 1.4 ± 0.39 | 1 ± 0.47 | NA | 1.3 ± 0.45 | NA | < 0.01<sup>2</sup> |
|  | median (min; max) | 2.4 (0.98; 4.2) | 1.4 (0.31; 2.6) | 0.91 (0.53; 1.9) | NA | 1.3 (0.38; 3.1) | NA |  |
| Serum NfL (pg/ml) | mean±SD | 12 ± 6.7 | 13 ± 7.2 | 18 ± 7 | 16 ± 7 | 15 ± 10 | 13 ± 6.7 | < 0.01<sup>2</sup> |
|  | median (min; max) | 10 (2.4; 51) | 11 (1.8; 77) | 14 (12; 29) | 14 (6.5; 39) | 13 (3.8; 81) | 11 (2; 63) |  |

<sup>1</sup> Pearson's Chi-squared test; <sup>2</sup> Kruskal-Wallis rank sum test
**Abbreviations:** Levodopa equivalent daily dosage (LEDD); education adjusted Montreal Cognitive Assessment (MoCA); Symbol Digit Modality Test (SDMT), executive function/working memory (WMS-III); Letter-Number Sequencing Test (LNS); Benton Judgment of Line Orientation test (BJLO); Movement Disorders Society Unified Parkinson's Disease rating Scale (MDS-UPDRS); Dopamine transporter imaging (DaT-Scan); Striatal Binding Ratio (SBR); Neurofilament light chain (NfL); Hopkins Verbal Learning Tests with immediate/total recall (HVLT-IR), discrimination recognition (HVLT-DG) and retention (HVLT-RT); Standard deviation (SD)

**Table 5:** Linear mixed effect model of log2NfL on diagnoses, time and their interactions, adjusted by age and sex

| Variable | Estimate | 2^Estimate | Std. Error | p-value |
| --- | --- | --- | --- | --- |
| (Intercept) | 0.5636 | 1.4780 | 0.1017 | **<0.0001** |
| Age | 0.0475 | 1.0335 | 0.0015 | **<0.0001** |
| Sex: male | -0.0947 | 0.9364 | 0.0299 | **0.0016** |
| Diagnostic Group: Genetic Cohort – Unaffected | 0.0774 | 1.0551 | 0.0494 | 0.1178 |
| Diagnostic Group: Prodromal | 0.1613 | 1.1183 | 0.0781 | **0.0392** |
| Diagnostic Group: Parkinson's Disease (PD) | 0.0615 | 1.0435 | 0.0466 | 0.1874 |
| Diagnostic Group: Genetic Cohort – PD | 0.1966 | 1.1460 | 0.0650 | **0.0025** |
| Diagnostic Group: Other Neurodegenerative Diseases (OND) | 0.5074 | 1.4215 | 0.1927 | **0.0086** |
| Time | 0.0008 | 1.0005 | 0.0094 | 0.9360 |
| Diagnostic Group: Genetic Cohort – Unaffected*TIME | 0.0039 | 1.0027 | 0.0175 | 0.8232 |
| Diagnostic Group: Prodromal*TIME | 0.0123 | 1.0085 | 0.0235 | 0.6022 |
| Diagnostic Group: Parkinson's Disease*TIME | 0.0541 | 1.0382 | 0.0111 | **<0.0001** |
| Diagnostic Group: Genetic Cohort – PD*TIME | 0.0358 | 1.0251 | 0.0150 | **0.0175** |
| Diagnostic Group: Other Neurodegenerative Diseases*TIME | 0.2165 | 1.1619 | 0.0435 | **<0.0001** |

With a linear median mixed model, when the adjusted NfL level doubled, the median MDS-UPDRS total score increased by 3.45 (FDR-adjusted p =0.0115); the median SDM total score decreased by 1.39 (FDR-p-value=0.026); the median HTLV-DG score decreased by 0.3 (FDR-adjusted p =0.03), and the median HTLV-RT score decreased by 0.029 (FDR-adjusted p=0.04). There was also a trend of increased MDS-UPDRS III with increased serum NfL (**table 6**).

**Table 6:** Associations between clinical measurements and age- and sex- adjusted log2NfL among patients with Parkinson’s disease in the PPMI validation cohort

| Variable | Estimate | Std. Error | Lower Bound | Upper Bound | p-value | FDR-adjusted p-value |
| --- | --- | --- | --- | --- | --- | --- |
| MDS-UPDRS part III | 1.3850 | 0.6214 | 0.1360 | 2.6340 | 0.0305 | 0.0731 |
| MDS-UPDRS total score | 3.4450 | 0.9802 | 1.4760 | 5.4150 | 0.0010 | **0.0115** |
| MoCA | -0.4153 | 0.2412 | -0.9001 | 0.0695 | 0.0915 | 0.1568 |
| HVLT-IR | -0.6134 | 0.3790 | -1.3750 | 0.1482 | 0.1120 | 0.1680 |
| HVLT-DG | -0.3019 | 0.1100 | -0.5230 | -0.0807 | 0.0085 | **0.0339** |
| HVLT-RT | -0.0286 | 0.0112 | -0.0512 | -0.0061 | 0.0138 | **0.0415** |
| SDMT | -1.3930 | 0.4657 | -2.3280 | -0.4567 | 0.0044 | **0.0261** |
| LNS | -0.0683 | 0.1339 | -0.3373 | 0.2007 | 0.6123 | 0.6679 |
| BJLOS | -0.2431 | 0.1404 | -0.5253 | 0.0390 | 0.0896 | 0.1568 |
| DaT-Scan PUTAMEN (SBR) | -0.0244 | 0.0253 | -0.0752 | 0.0265 | 0.3398 | 0.4531 |
| DaT-Scan CAUDATE (SBR) | -0.0225 | 0.0420 | -0.1069 | 0.0618 | 0.5937 | 0.6679 |
| DaT-Scan STRIATUM (SBR) | 0.0038 | 0.0400 | -0.0766 | 0.0841 | 0.9254 | 0.9254 |

**Abbreviations:** Education adjusted Montreal Cognitive Assessment (MoCA); Symbol Digit Modality Test (SDMT), Letter-Number Sequencing Test (LNS); Benton Judgment of Line Orientation test (BJLO); Movement Disorders Society Unified Parkinson's Disease rating Scale (MDS-UPDRS); Dopamine transporter imaging (DaT-Scan); Striatal Binding Ratio (SBR); Hopkins Verbal Learning Tests with immediate/total recall (HVLT-IR), discrimination recognition (HVLT-DG) and retention (HVLT-RT)

## Discussion

We investigated NfL levels by sandwich immunoassays in longitudinal and cross-sectional discovery cohorts from a single center and validated the findings in a longitudinal multicenter cohort consisting of HC, prodromal, and established PD participants. We also studied symptomatic and asymptomatic mutation carriers of known PD genetic mutations with follow-ups of up to six years.

Results obtained from our discovery cohorts indicated that serum and CSF NfL levels were highest in OND and higher in PD participants compared with controls; CSF NfL levels in PD increased non significantly longitudinally over four years.

In the second step, we focused on NfL serum levels in the validation cohort, considering the strong correlation between CSF and serum NfL levels and the advantage of using a peripheral and less invasive source for the specimen. The main findings in the validation cohort were: (1) mean levels of serum NfL were higher in established PD versus HC at each of the six time points during the first five years after diagnosis; (2) markedly higher levels of NfL were seen in the few participants with OND, including participants with DLB and MSA (discussed in more detail below) (3) there was a significant (age-adjusted) longitudinal increase in serum NfL in PD compared to HC and (4) the longitudinal change in serum NfL correlated significantly with MDS-UPDRS total scores and some cognitive measures, suggesting that serum NfL may be a biomarker of clinical progression (independent of age and sex). Overall, the longitudinal analysis of serum NfL in the validation set confirmed the findings from the discovery cohorts. The PPMI cohort enabled us to analyze prodromal conditions and a genetic cohort with symptomatic and asymptomatic mutation carriers. We found that the serum NfL levels of participants with two prodromal conditions for *α*-synuclein aggregation disorders had higher levels than PD or HC and lower than OND, including MSA and DLB. This was as expected because iRBD participants especially have a high risk to predominantly convert into PD but also to DLB and MSA. The 17 MSA cases in the OND group of the cross-sectional discovery cohort showed the highest median values in CSF [2829.86 pg/ml; range: 794.32, 6015.71] and in serum [37.42 pg/ml; range: 21.26, 91.99]. In the genetic cohort, serum NfL levels of asymptomatic mutation carriers in all three mutation carrier groups were lower than in their respective symptomatic group. The levels in all six genetic groups (symptomatic and asymptomatic) remained relatively stable over the two to three years of follow-up and there were no differences in NfL rate of change between the mutation carrier groups. Subsequent analyses will explore the effects of disease duration on changes in serum NfL.

Elevated levels of NfL, as seen here in PD and OND compared with HC, have been identified in several other neurological conditions including dementia disorders and multiple sclerosis. Therefore, this marker for axonal damage is not specific for any disease,^4^ but could be useful for exploring specific questions within disease entities. Within PD spectrum disorders, NfL may be particularly useful in discriminating PD from cognate disorders such as MSA, PSP, and DLB, as has been previously described.^7^ Due to the high rate of misdiagnoses in early PD as shown in a neuropathologically-confirmed cohort,^2^ there is a need for biomarkers to distinguish sporadic PD from other neurodegenerative disorders with Parkinson syndromes, such as the two -synuclein aggregation disorders, DLB with early dementia and marked hallucinations and MSA with clinically severe autonomic disturbances and -synuclein deposition in the glia (and not in neurons), thus, disorders with a completely different clinical and neuropathological disease phenotype. This is important in clinical practice as “Parkinson-plus” syndromes may have a completely different underlying pathophysiology, have worse prognoses and may ultimately require different therapy regimens. This need will grow as medicine becomes more personalized. In addition, the ability to select patients for inclusion or stratification within clinical trials is also becoming increasingly important. The diagnosis of PD in PPMI is clinical; it is based on systematic history and examination, structural MRI (to rule out other diseases), pathological DaT by central read, and most importantly by longitudinal clinical follow-up. Based on this process, eight participants in the PD cohort analyzed here were found to have diagnoses other than PD after five years follow-up and were thus taken out of the PD cohort. This group of participants, despite the small size, was separated in the analysis as OND. Serum NfL levels in this OND group were higher at baseline (before the follow-up and before evolving into another disease) and may provide future diagnostic utility. Additional studies including analysis on samples from post-mortem PD-confirmed participants will further establish the utility of serum NfL measurements in differentiating PD from OND. In numerous previous CSF studies (conducted before ultrasensitive technologies were available), the mean NfL levels in OND were found to be markedly increased compared to HC, such as in multiple sclerosis (4.5-fold increase), traumatic brain injury (3 fold increase), PSP, corticobasal degeneration, MSA (3-4.25 fold increase) as well as in other, more slowly progressing neurodegenerative disorders such as Alzheimer’s disease (1.5-2-fold increase).^23^ Compared to these disorders, the increase is smaller in PD (1.25 in CSF and 1.4 in serum), where *α*-synuclein aggregation occurs mainly in neurons with high energy turnover in less myelinated axons,^24^ while NfL is mainly expressed in larger myelinated axons.^25^ Also, in contrast to many more rapidly progressive or acute diseases, the pathological changes in PD and the respective cell loss is only mild.^26^

Despite the slight increase shown here, the relatively higher serum NfL levels in the prodromal groups (compared to PD and HC), especially in the iRBD group, may indicate the presence of active disease and potential for conversion to either PD or Parkinsonian syndromes. Furthermore, longitudinal NfL levels may, in fact, be highest in the early stages of the disease with the greatest disease activity as has been similarly seen in *β*-amyloid in Alzheimers disease.^27^ NfL levels are also higher in presymptomatic cases with amyotrophic lateral sclerosis. ^28^

Another variable affecting the increase of NfL levels is age. Across published studies, NfL in CSF and blood increases with age,^23^ as we also identified in the PPMI cohort. For example, the prodromal groups in PPMI are on average 3-5 years older than the PD group and the iRBD participants are older than the hyposmic participants featuring higher NfL values. The reasons for this positive association of serum NfL and age is explained by structural alterations of the axons with ageing, including vascular disease, metabolic changes, and inflammation. All of these have also been shown to play a role in PD progression^29^, which may also influence the NfL levels in blood. To account for these systematic differences at baseline we adjusted all our serum analyses for age and sex.

Similar to a previous report showing that CSF NfL levels were relatively stable despite disease progression in PD over 12 months^9^ we have observed a slight increase of serum NfL over 72 months follow-up, which did not correlate with motor progression. The discrepancy between CSF and serum measurement could possibly be due to the smaller CSF sample size. In contrast, NfL levels in serum increased significantly over 60 months follow-up in the validation cohorts. The longitudinal increase in the age- and sex-adjusted serum NfL significantly correlated with changes over time in MDS-UPDRS total scores in the PPMI cohort. A significant association with longitudinal progression of age- and sex-adjusted serum NfL was also seen for the Hopkins Verbal Learning Test discrimination recognition and retention as well as for processing speed/attention test SDMT.

We did not observe a longitudinal change of serum NfL in the prodromal cohort. This might be due to small sample size and the heterogeneity of the group as recruited, both in terms of disease stage as well as eventual diagnostic category and phenotypes of individual participants. The apparent stability over time in the prodromal cohort could also indicate that serum NfL levels do not change continuously before motor symptoms evolve. This suggestion is also supported by the longitudinal stability of serum NfL in the asymptomatic mutation carriers. Only continued follow-up will identify participants converting to a motor disease or symptomatic disease state. The individual levels may indicate the prognostic direction.

We identified NfL as the first blood-based PD progression biomarker. We are aware that the diagnostic accuracy of NfL in CSF and serum for PD vs. HC is small (with areas under the receiver operating curves around 0.6). Nevertheless, we observed slightly higher levels of serum NfL in PD compared to HC even at early stages of the disease, a mild longitudinal increase in serum NfL levels over time and correlations of serum NfL with clinical measures of disease progression in PPMI (but not in CSF NfL in DeNoPa). Increases in serum NfL were, in general, less than those seen in OND, which are either more rapidly progressive than PD, associated with damage to myelinated tracts (as seen in multiple sclerosis), or associated with significantly greater cell death (as seen in Alzheimer’s and extreme in Creutzfeldt-Jakob disease). Our NfL data will be strengthened with continued analyses of these data using additional alternative statistical models as well as follow-up of the cohorts, especially in participants at risk for disease progression. In addition, monitoring of larger longitudinal cohorts with a focus on prodromal or asymptomatic PD with longer observational time to allow the development of motor disease are needed and will be pursued in the extended PPMI 2.0 starting 2020. This being said, we remain cognizant that increased NfL levels are not specific to PD or any other neurodegenerative disorder. Thus, more specific markers will need to be identified, leading hopefully to a panel of different markers reflecting disease state, rate, and fate. Finally, we note the profound influence of age and sex on serum NfL levels. We, therefore, recommend that age- and sex-based adjustments be applied when interpreting serum NfL levels in clinical research and practice.

## Acknowledgements

We thank the Michael J. Fox Foundation, all of our PPMI colleagues and the many individuals who have given their time and of themselves to be participants in this study. This study was funded by The Michael J. Fox Foundation for Parkinson’s Research and funding partners including Abbvie, Allergan, Avid Radiopharmaceuticals, Abbott, Biogen Inc, BioLegend, Bristol-Myers Squibb, Celgene, Denali, GE Healthcare, Genentech, GlaxoSmithKline, Lilly, Lundbeck, Merck, Meso Scale Discovery, Pfizer, Piramal, Prevail Therapeutics, F. Hoffman-La Roche Ltd., Sanofi Genzyme, Servier, Takeda, Teva, UCB.

## Author contributions

BM, DG, and JMC conceived the study. BM, SH, and DG designed the study and were responsible for data processing; MD, FG, WW, RG, DG and BM oversaw all statistical analyses; NK, HZ and DG were involved in sample analyses and data interpretation; MF, KM, LMC, SH and AS oversaw patient recruitment and assisted in the interpretation of data. BM wrote the manuscript. MD, WW, FG, DG, TF, HZ, SS, RG, NK, MF, LMC, TS, ABS, DW, KM, AS, JMC, SH, DG, CMT and CT co-edited the manuscript. BM, SH, DG, and MD had full access to the clinical primary data and take responsibility for the integrity of the data and the accuracy of the data analysis. All authors had access to the data generated in the study including the statistical analysis and decided to submit the paper for publication.

## Declaration of interests: financial disclosures of all authors (for the preceding 3 years)

Brit Mollenhauer has received honoraria for consultancy from Roche, Biogen, UCB, and Sun Pharma Advanced Research Company and has received research funding from the Deutsche Forschungsgemeinschaft (DFG), EU (Horizon2020), Parkinson Fonds Deutschland, Deutsche Parkinson Vereinigung and the Michael J. Fox Foundation for Parkinson’s Research.

Mark Frasier and Samantha Hutten are employees of the Michael J. Fox Foundation for Parkinson’s Research.

Andrew Siderowf has received honoraria for consultancy from Biogen, Voyager Therapeutics, Merck, and Denali; honoraria for data and safety monitoring from Wave Life Sciences, Prilenia Therapeutics; and grant funding from the Michael J. Fox Foundation for Parkinson’s Research and NINDS.

John Seybl is a consultant for Roche, GE Healthcare, Biogen, Life Medical Imaging, Invicro, LikeMinds. Equity interest: Invicro

Caroline M Tanner is an employee of the University of California – San Francisco and the San Francisco Veterans Affairs Health Care System. She receives grants from the Michael J. Fox Foundation, the Parkinson’s Foundation, the Department of Defense, BioElectron, Roche/Genentech, Biogen Idec and the National Institutes of Health, compensation for serving on Data Monitoring Committees from Biotie Therapeutics, Voyager Therapeutics and Intec Pharma and personal fees for consulting from Neurocrine Biosciences, Adamas Therapeutics, Biogen Idec, 23andMe, Alexza, Grey Matter, Acorda, Acadia and CNS Ratings.

Lana M. Chahine receives research support from the Michael J Fox Foundation, has received travel payment from MJFF to MJFF conferences, is a paid consultant to MJFF, receives research support from the UPMC Competitive Medical Research Fund, is study site investigator for a study sponsored by Biogen, is a site sub-investigator for a study sponsored by Voyager, received payment from Elsevier (for book authorship), and receives royalties from Wolters Kluwel (for book authorship).

Kenneth Marek has served as a consultant for Pfizer, GE Healthcare, Merck, Lilly, BMS, Piramal, Prothena, Neurophage, nLife, Roche, and receives funding for the following grants: W81XWH-06-1-0678 Establishing an ‘at risk’ cohort for Parkinson Disease Neuroprevention using olfactory testing and DAT imaging, DOD, Investigator 10/1/06 – 09/30/15; Parkinson Progression Marker Initiative (PPMI), Michael J. Fox Foundation, Principal Investigator 6/15/09 – 6/14/18; DAT imaging in LRRK2 family members, the Michael J. Fox Foundation, Principal Investigator 1/15/10 – 1/14/15. Ownership in Molecular NeuroImaging,LLC.

Douglas Galasko is supported by NIH grant AGO5131, and by the Michael J. Fox Foundation. He has provided consultation for vTv Pharmaceuticals, Eli Lilly, Inc, and Proclara, Inc.

Karl Kieburtz serves as Consultant for Clintrex LLC, Roche/Genentech, Novartis, Blackfynn and has grant support from NIH (NINDS, NCATS), Michael J Fox Foundation; Ownership: Clintrex LLC, Hoover Brown LLC, Safe Therapeutics LLC

Tanya Simuni has served as a consultant for Acadia, Abbvie, Adamas, Anavex, Aptinyx, Allergan, Accorda, Denali, Neuroderm, Neurocrine, Revance, Sanofi, Sunovion, TEVA, Takeda, Voyager, US World Meds, Parkinson’s Foundation, and the Michael J. Fox Foundation for Parkinson’s Research; Dr. Simuni has served as a speaker and received an honorarium from Acadia, Adamas, and TEVA; Dr Simuni is on the Scientific advisory board for Neuroderm, Sanofi, and MJFF. Dr. Simuni has received research funding from the NINDS, Parkinson’s Foundation, MJFF, Biogen, Roche, Neuroderm, Sanofi, Sun Pharma.

Tatiana Foroud is supported by NIH and the Michael J. Fox Foundation,

Jesse Cedarbaum is a former employee of and shareholder in Biogen.

TYL, WW, DG and FG are employees of Biogen

SS has received salaries from the EU Horizon 2020 research and innovation program under grant agreement No. 634821 and from the Deutsche Forschungsgemeinschaft (DFG) under grant agreement No. MO 2088/5-1.

HZ has participated in scientific advisory board meetings for Roche Diagnostics, Wave, Samumed and CogRx, has given lectures in symposia sponsored by Biogen and Alzecure and Fujirebio, and is a co-founder of Brain Biomarker Solutions in Gothenburg AB, a GU Ventures-based platform company at the University of Gothenburg (all outside submitted work).

CT has received grants from M.J. Fox, European grant: program Horizon 2020 and consulting fees from Britannia, Roche as well as honoraria from UCB, Gruenenthal, Otsuka. Book for PD patients (Schattauer publisher), PDSS2 Scale, European patent on Dyskinesias of PD: Commercial interest: no

Dr. Weintraub has received research funding or support from Michael J. Fox Foundation for Parkinson’s Research, Alzheimer’s Therapeutic Research Initiative (ATRI), Alzheimer’s Disease Cooperative Study (ADCS), the International Parkinson and Movement Disorder Society (IPMDS); honoraria for consultancy from Acadia, Aptinyx, Biogen, CHDI Foundation, Clintrex LLC, Enterin, F. Hoffmann-La Roche Ltd, Ferring, Promentis, Signant Health, Sunovion, and Takeda; and license fee payments from the University of Pennsylvania for the QUIP and QUIP-RS.

AWT, RG, MD, ABS and NK have no disclosures to report.

